# Multimodal small-molecule screening for human prion protein binders

**DOI:** 10.1101/2020.06.18.159418

**Authors:** Andrew G Reidenbach, Michael F Mesleh, Dominick Casalena, Sonia M Vallabh, Jayme L Dahlin, Alison J Leed, Alix I Chan, Dmitry L Usanov, Jenna B Yehl, Christopher T Lemke, Arthur J Campbell, Rishi N Shah, Om K Shrestha, Joshua R Sacher, Victor L Rangel, Jamie A Moroco, Murugappan Sathappa, Maria Cristina Nonato, Kong T Nguyen, S Kirk Wright, David R Liu, Florence F Wagner, Virendar K Kaushik, Douglas S Auld, Stuart L Schreiber, Eric Vallabh Minikel

## Abstract

Prion disease is a rapidly progressive neurodegenerative disorder caused by misfolding and aggregation of the prion protein (PrP), and there are currently no therapeutic options. PrP ligands could theoretically antagonize prion formation by protecting the native protein from misfolding or by targeting it for degradation, but no validated small-molecule binders have been discovered to date. We deployed a variety of screening methods in an effort to discover binders of PrP, including ^19^F-observed and saturation transfer difference (STD) nuclear magnetic resonance (NMR) spectroscopy, differential scanning fluorimetry (DSF), DNA-encoded library selection, and *in silico* screening. A single benzimidazole compound was confirmed in concentration-response, but affinity was very weak (*K_d_* > 1 mM), and it could not be advanced further. The exceptionally low hit rate observed here suggests that PrP is a difficult target for small-molecule binders. While orthogonal binder discovery methods could yield high affinity compounds, non-small-molecule modalities may offer independent paths forward against prion disease.

## INTRODUCTION

Prion disease is a rapidly progressive neurodegenerative disorder caused by misfolding and aggregation of the prion protein, or PrP^1^. No effective therapeutics currently exist for prion disease, but PrP is a genetically and pharmacologically validated drug target^2^. PrP-lowering antisense oligonucleotides (ASOs) are in preclinical development^3–5^, and PrP-binding antibodies have been tested preclinically^6^ as well as clinically in a compassionate use context^7^. Here, we sought to augment the therapeutic pipeline by discovering small molecules that bind PrP.

In principle, small molecules could prevent or treat prion disease by protecting PrP from misfolding or by lowering its abundance. By sterically blocking interactions with misfolded PrP, or simply through the free energy of binding, a chaperone might stabilize PrP against misfolding, following precedents in transthyretin amyloidosis^8,9^ and cystic fibrosis^10,11^. Proofs of principle for this approach include the efficacy of monoclonal antibodies to PrP to clear prion infection in cell culture^12,13^ and in peripheral tissues of animals^14^ as well as the stability of PrP “stapled” with non-native disulfide bonds^15^. Alternatively, small-molecule binding events can sometimes directly lead to protein degradation^16,17^, and if not, a binder could serve as a starting point for engineering a bifunctional molecule to specifically target PrP for degradation^18,19^. Although at present most bifunctional degrader strategies are best suited to intracellular targets owing to reliance on cytoplasmic E3 ubiquitin ligases, recent studies suggest alternate routes to targeted degradation of cell surface proteins^20^ such as PrP.

Decades of effort have not yet yielded a small-molecule PrP binder suitable for advancement as a drug candidate^21^. The development of phenotypic screening for antagonists of misfolded PrP accumulation in cultured cells^22^ enabled the identification of several compounds effective *in vivo*^23–26^, but advancement of these compounds has been hindered by lack of activity against human prion strains and unclear mechanisms of action^26–29^. Meanwhile, several compounds shown to interact with PrP through biophysical assays have demonstrated antiprion activity in a range of experimental systems^30–34^. However, these compounds likewise appear to lack clinical promise as none are simultaneously specific^35^, potent, and drug-like. Certain metallated porphyrins^36,37^ interact with PrP with affinity values comparable to their effective concentrations in cell culture^36,37^, and some exhibit *in vivo* activity in certain paradigms^38^. Similarly, a range of anionic polymers^33,39–42^ also bind PrP and show *in vivo* antiprion activity in certain contexts^41,43,44^. However, these binding events may not be monomeric^45^ nor specific to PrP^35,46^. Still other compounds with demonstrated antiprion activity exhibit interaction with PrP only at concentrations orders of magnitude above their effective concentration in cell culture^47,48^.

Because PrP’s biology does not lend itself to enzymatic or activity assays, we chose to apply several non-activity-based screening modalities: ^19^F-observed and saturation transfer difference (STD) NMR fragment screening, differential scanning fluorimetry (DSF), DNA-encoded library (DEL) selection, and an *in silico* screen. We selected a fragment-based drug discovery paradigm as a starting point based on PrP’s small size and lack of obvious binding pockets^49^, and the success of this method in identifying ligands for targets refractory to other approaches^50^. Our campaign utilized ^19^F NMR and STD NMR because these approaches are sensitive to weak affinity binders^51^, have low false positive rates^52^ and allow searching a large swath of chemical space through small, highly soluble fragments that can later be optimized into larger, higher affinity molecules^53^. DSF was employed to find compounds that directly influence the thermal stability of PrP. This technique quantifies protein thermal stability by measuring fluorescence of a solvatochromic dye (SYPRO Orange) as a function of temperature as it binds to unfolded regions of a protein^54^. In principle, DSF hits should have the desired property of stabilizing the target protein. DNA-encoded library (DEL) selection uses a pool of thousands to trillions of individually DNA-barcoded molecules that are added to an immobilized recombinant protein. Nonspecific molecules are washed away, and putative binders are eluted, PCR amplified, and subjected to next-generation DNA sequencing for identification. We used a DEL of peptide macrocycles^55^, which display better stability and have a lower entropic cost of binding compared to linear peptides, making them suitable for targeting surfaces of proteins^56,57^. Finally, we employed the artificial intelligence-based *in silico* screening method, AtomNet^®58^, which uses a protein structure-based, convolutional neural network to predict molecular binding affinities. This technique was recently used to discover a selective binder and degrader of Miro1^59^.

## RESULTS

### Fragment-based drug discovery through NMR screening

From five commercial and internal sources (Figure 1A) we selected 6,630 low molecular weight, high solubility fragments for a fragment-based drug discovery campaign. The compounds in these libraries were mostly small (<300 Da) and had a range of cLogP values and hydrogen bond donor and acceptor sites that mostly fell within the Rule of Three^60^ for fragment-based screening (Figures 1B-D). Fragments were screened against either HuPrP23-231 or HuPrP90-231 using pooled, ^19^F ligand-observed or STD NMR methods as a primary screen, singleton STD NMR for re-testing, and protein-observed ^1^H-^15^N TROSY NMR for validation (Table 1).

**Figure 1.**
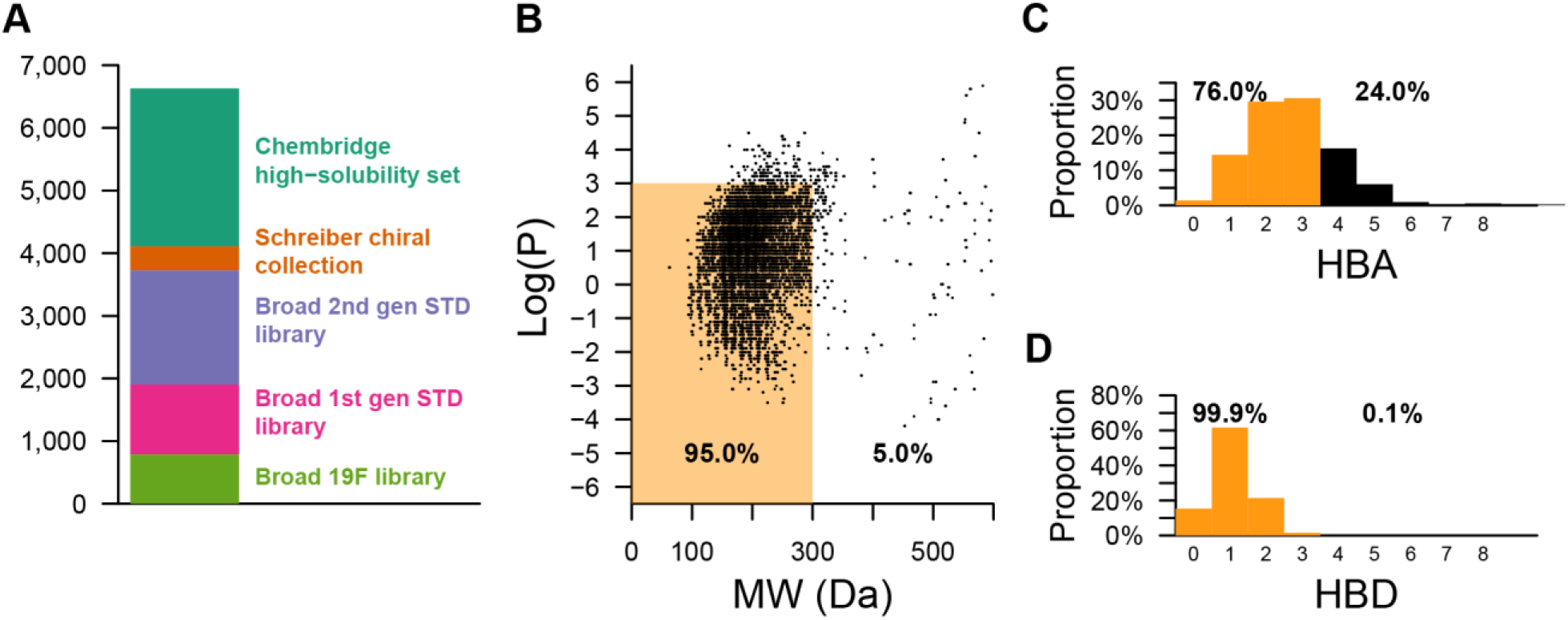
Physicochemical properties of fragment libraries screened. A) Composition of fragment libraries. B) Scatter plot of log(P) versus molecular weight (MW) for all fragments screened against PrP using ^19^F or STD NMR. C and D) Number of hydrogen bond donors (HBD) or hydrogen bond acceptors (HBA). Orange denotes compliance with the “Rule of Three”^60^ and black denotes noncompliance.

**Table 1.** Summary of NMR fragment screening. “Compounds screened” lists the total number of compounds in a given collection of molecules. “Pools with hits” indicates the number compound pools with observed hits. “Retested by STD” indicates the number of individual compounds that were retested for STD signal from each hit pool. “Retested by TROSY” shows the number of compounds that gave an STD signal and were therefore advanced to 2D TROSY NMR.

| Library name | Compounds screened | Pools with hits | Retested by STD | Retested by TROSY | Validated hits |
| --- | --- | --- | --- | --- | --- |
| Broad Institute $^{19}\text{F}$ library | 785 | 14 | 14 | 5 | 0 |
| Broad Institute 1st gen STD library | 1,116 | 16 | 15 | 4 | 0 |
| Broad Institute 2nd gen STD library | 1,823 | 43 | 55 | 34 | 1 |
| Schreiber chiral collection | 381 | 1 | 5 | 0 | 0 |
| ChemBridge High Solubility Subset | 2,525 | 31 | 149 | 37 | 0 |
| Total | 6,630 | 105 | 238 | 80 | 1 |

Of 6,630 compounds tested, 238 initial hits were re-tested as singletons by STD NMR, of which 80 were further tested by TROSY NMR. A single compound 5,6-dichloro-2-methyl-1*H*-benzimidazole (**1**) gave a reproducible STD signal in the presence of PrP (Figure 2A) and induced chemical shift perturbations (CSPs) in the TROSY spectrum of HuPrP90-231 (Figures 2B-C, S1, and S2A). Mapping these residues onto an NMR structure of HuPrP (PDB 1HJM)^61^ revealed no discernable pocket, with shifts scattered across the structure (Figure 2D). Nonetheless, we observed similar resonance shifts in the full-length protein HuPrP23-231, suggesting that this binding is not an artifact of using a truncated construct (Figure S2A), and CSPs were confirmed to be dose-responsive for several residues (Figures 2E and S2B), and the compound caused a small (~0.2°C) but significant decrease in melting temperature by DSF (Figure S2D).

**Figure 2.**
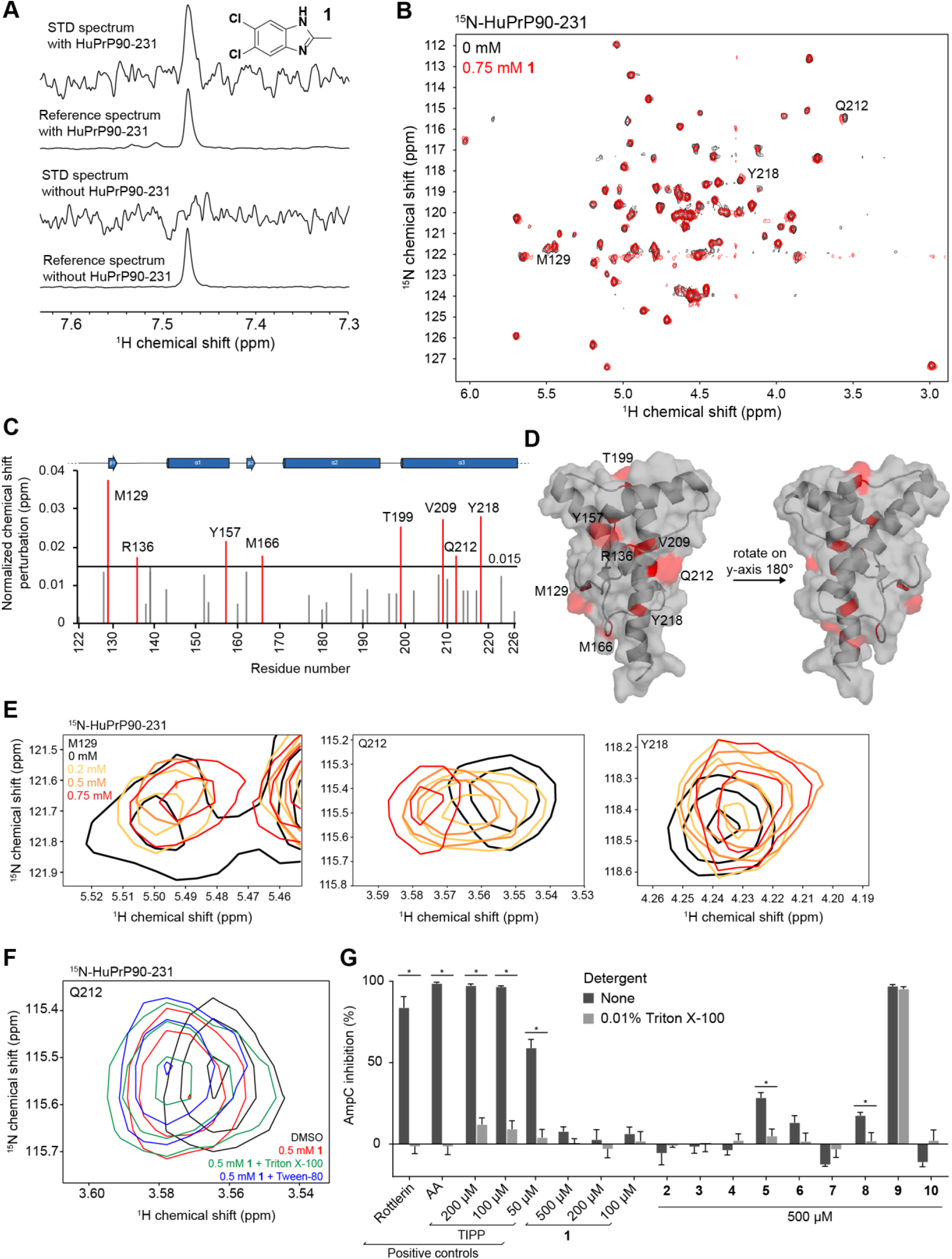
Validation and characterization of a benzimidazole fragment hit. A) STD NMR spectra of **1** (5,6-dichloro-2-methyl-1H-benzimidazole) with and without HuPrP90-231. STD spectra are scaled to 16X the reference spectra. B) TROSY spectrum of ^15^N-HuPrP90-231 with DMSO (black) or 0.75 mM **1** (red). Peaks that shift greater than 0.015 ppm are denoted with the residue number. C) Normalized chemical shift perturbations upon addition of 0.75 mM compound **1**. D) Residues that shift more than 0.015 ppm were mapped onto the NMR structure of PrP (PDB 1HJM)^61^. E) Concentration dependent CSPs of residues Q212, M129, and Y218 upon addition of **1**. F) ^1^H-^15^N TROSY chemical shifts in the presence of 0.75 mM **1** with and without detergents. G) AmpC inhibition assay of the **1** and its analogs. Rottlerin (10 μM), anacardic acid (AA, 10 μM), and tetraiodophenolphthalein (TIPP) are prototypical aggregators. Adding detergent to small-molecule aggregates dissociates them and attenuates inhibition of AmpC. *, significance cutoff between detergent and non-detergent tests was p & 0.01 after correction for multiple comparisons, data are mean ± SD of four intra-run technical replicates performed on same microplate.

Because the CSPs caused by **1** are so small, we wanted to be certain that this compound was not perturbing HuPrP due to nonspecific colloidal aggregation. To test whether **1** is causing CSPs due to aggregation, ^15^N-HuPrP90-231 was incubated with **1** in the presence or absence of nonionic detergents Triton X-100 or Tween-20. The CSPs resulting from **1** were preserved in the presence of detergent, suggesting that **1** is not an aggregator (Figure 2F). To assess compound aggregation by an orthogonal method, we used the well-established AmpC β-lactamase inhibition assay^62^. AmpC is inhibited by small molecules that form aggregates, and these aggregates can be disrupted by addition of detergent. No significant inhibition of AmpC was observed with **1** even at 500 μM concentrations, while the positive control compounds rottlerin and anacardic acid (AA) showed inhibition at 10 μM that could be relieved upon detergent addition (Figure 2G). Analogs of **1** (compounds **2-10**) were also tested; the majority of them did not inhibit AmpC, and none inhibited AmpC as well as the positive controls. Collectively, these data argue that **1** is not an aggregator and does not cause PrP CSPs via aggregation.

The small magnitude of CSPs and lack of saturation at concentrations up to 0.75 mM suggested that **1** has a *K*_d_ for PrP in the millimolar range, too weak to interrogate by many non-NMR orthogonal biophysical assays. In an attempt to find a stronger binder, we tested 54 analogs of **1** from commercial sources and the Broad Institute’s internal library by STD NMR and TROSY (Table 2, Table S1, and Figure S3). Of the 20 compounds most similar to **1**, 11 demonstrated positive STD and TROSY signal (Table 2), however, none of the analogs had TROSY CSPs larger than **1** by visual inspection, and thus they were not subjected to further biophysical assays. Despite efforts to soak unliganded PrP crystals with **1** and 20 of its analogs, no electron density attributable to a compound was identified (Table S1).

**Table 2.**
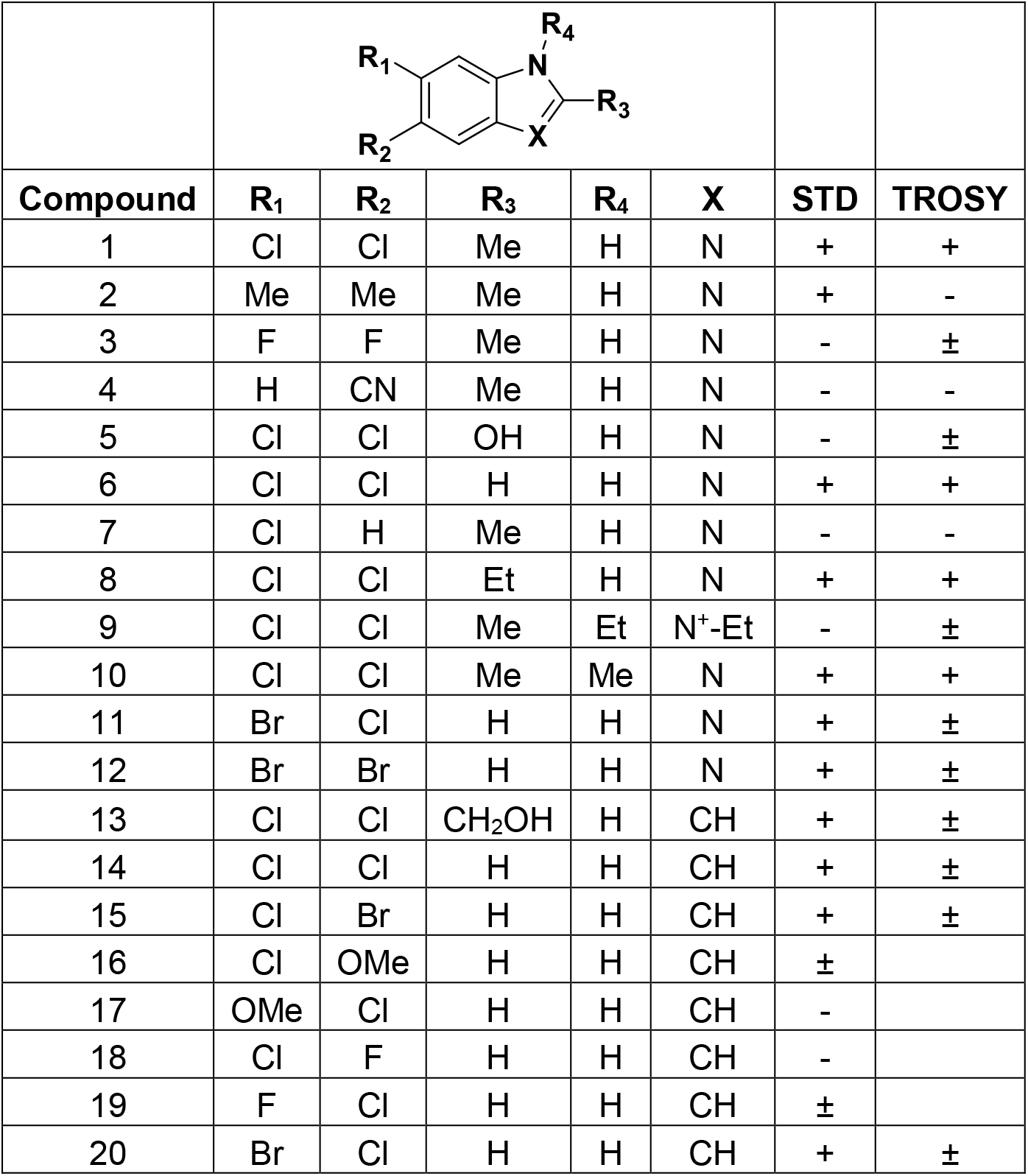
Analogs of compound 1 tested by STD and TROSY NMR. Analogs of **1** were initially tested by STD NMR and positive STD hits were assayed by TROSY. + indicates positive STD or TROSY signal; - indicates no STD or TROSY signal; ± indicates borderline positive signal. Blank cells indicate that the analog was not tested. All spectra were assessed by visual inspection.

|  |  |  |  |  |  |  |  |
| --- | --- | --- | --- | --- | --- | --- | --- |
| Compound | R<sub>1</sub> | R<sub>2</sub> | R<sub>3</sub> | R<sub>4</sub> | X | STD | TROSY |
| 1 | Cl | Cl | Me | H | N | + | + |
| 2 | Me | Me | Me | H | N | + | - |
| 3 | F | F | Me | H | N | - | ± |
| 4 | H | CN | Me | H | N | - | - |
| 5 | Cl | Cl | OH | H | N | - | ± |
| 6 | Cl | Cl | H | H | N | + | + |
| 7 | Cl | H | Me | H | N | - | - |
| 8 | Cl | Cl | Et | H | N | + | + |
| 9 | Cl | Cl | Me | Et | N<sup>+</sup>-Et | - | ± |
| 10 | Cl | Cl | Me | Me | N | + | + |
| 11 | Br | Cl | H | H | N | + | ± |
| 12 | Br | Br | H | H | N | + | ± |
| 13 | Cl | Cl | CH<sub>2</sub>OH | H | CH | + | ± |
| 14 | Cl | Cl | H | H | CH | + | ± |
| 15 | Cl | Br | H | H | CH | + | ± |
| 16 | Cl | OMe | H | H | CH | ± |  |
| 17 | OMe | Cl | H | H | CH | - |  |
| 18 | Cl | F | H | H | CH | - |  |
| 19 | F | Cl | H | H | CH | ± |  |
| 20 | Br | Cl | H | H | CH | + | ± |

### Thermal shift screening

We tested a library of 30,013 compounds from the Novartis Screening Set for External Collaborations (SSEC) in singleton for thermal stabilization of PrP using DSF (Figures 3 and S5A-B). An ideal DSF screen would possess a high signal-to-baseline fluorescence ratio, tightly distributed melting temperatures in the apo condition, and a ≥10:1 molar ratio of soluble ligand to protein^54^. We varied a range of assay parameters including protein concentration, dye concentration, assay volume, use of HuPrP90-231 or HuPrP23-231, and buffer conditions including buffering agent, metals, and DMSO concentration (Figure S4). We obtained acceptable melt curves only at high protein concentrations (Figure S4). Our final screening conditions achieved a ~5:1 signal-to-baseline fluorescence and a 0.06 °C median absolute deviation (MAD) with 30 μM HuPrP90-231; median T_m_ was 68.5 °C (Figure 3A). Compounds were screened at 100 μM for a >3:1 ligand-to-protein ratio, though we lack empirical data on their solubility over the temperature ramp. We chose hit compounds that either positively or negatively affected the melting temperature (T_m_) of PrP based on separate criteria (see Methods). An internally developed pipeline was used to perform Boltzmann fitting of the fluorescence data and call T_m_ values. Irregular melt curves were automatically flagged and discarded. Two hit criteria were chosen for positive T_m_ shifters: 1) a statistical cutoff of greater than 3*MAD of the DMSO control wells (0.17 °C) and 2) an initial fluorescence of less than 6 to eliminate compounds that have intrinsic fluorescence or distort the melt curve. Because there were so many negative shifters, stricter hit cutoffs of −0.7 °C > ΔT_m_ > −9 °C were applied. Even though negative shifters are predicted to destabilize PrP, one hypothesis is that such compounds bind a partially folded or destabilized form of PrP and could exhibit antiprion properties^63^. Compounds that passed our hit criteria were retested by DSF in triplicate (Figure 3C). Here, we applied stricter hit cutoffs (see Methods) due to throughput limitations of our orthogonal heteronuclear single quantum coherence spectroscopy (HSQC) NMR assay. The 183 reproducible positive hits were passed through frequent hitter and PAINS filtering^64^, narrowing the list to 117 compounds, of which we were able to test 93 for PrP binding by HSQC using ^15^N-HuPrP90-231. Even though the melt curves of PrP with these compounds were often very robust and reproducible (Figure 3D), none of the compounds tested by HSQC led to PrP CSPs at 100 μM concentration (Figures 3E-F). This suggested that the observed thermal shifts were not mediated by binding PrP, and were likely artifacts. In support of this interpretation, when we tested eight of the validation screen hits by an orthogonal thermal shift method, differential scanning calorimetry, we were unable to reproduce the change in melting temperature (ΔT_m_) observed in DSF (Figures S5D-E).

**Figure 3.**
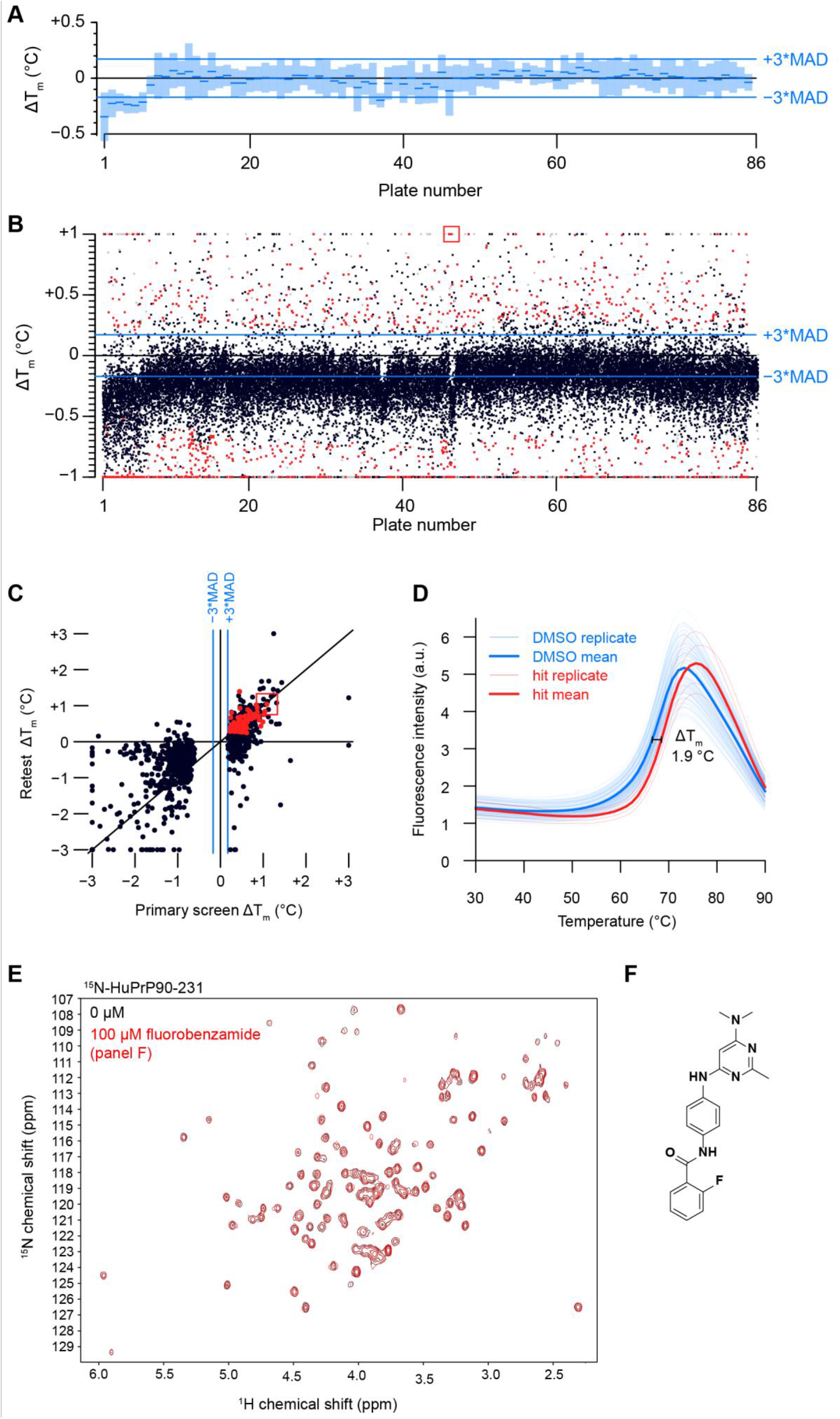
Thermal shift screening. A) A box plot of the HuPrP90-231 median T_m_ (blue dash) for the DMSO controls from each of the 86 384-well plates screened. These values are relative to the median from all plates combined. Light blue rectangles around each median represent ±3*MAD. B) Scatter plot of ΔT_m_ data from the initial screen with red dots indicating hits that were screened in triplicate. Red dots were compounds called as hits (see Methods for details). Grey dots were compounds that were flagged for having melt curve analysis errors. Black dots are compounds that did not meet our hit calling threshold but were not flagged. Compounds that resulted in PrP shifts of greater than or less than one degree were plotted at +1 or −1, respectively. C) Scatter plot of ΔT_m_ values from primary screening versus triplicate screening data. D) Fluorescence melt curves of HuPrP90-231 with DMSO and a positive ΔT_m_ shifter (fluorobenzamide shown in panel F). n = 128 for DMSO and n = 4 for test compound. E) HSQC spectrum of ^15^N-HuPrP90-231 with hit (fluorobenzamide shown in panel F; 100 μM). F) Chemical structure of selected fluorobenzamide hit, boxed in red in panels B and C.

**Table 3.**
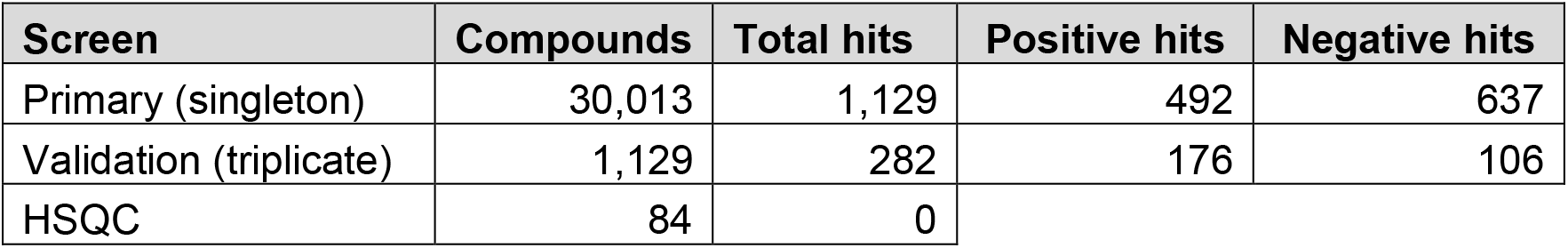
Summary of thermal shift screening results. “Compounds” provides the total number of compounds tested in that screening step. “Positive hits” and “Negative hits” list the number of molecules that shifted the T_m_ of HuPrP90-231 positively or negatively, respectively.

| Screen | Compounds | Total hits | Positive hits | Negative hits |
| --- | --- | --- | --- | --- |
| Primary (singleton) | 30,013 | 1,129 | 492 | 637 |
| Validation (triplicate) | 1,129 | 282 | 176 | 106 |
| HSQC | 84 | 0 |  |  |

### DNA-encoded library selection

We performed a selection using HuPrP90-231 with a DEL library of 256,000 macrocycles. Barcode rank abundance in the unenriched library was plotted against enrichment observed in the PrP condition versus a no-protein control condition, revealing enriched compounds across three structural scaffolds (Figure 4A). The *KRD scaffold was judged to be a likely covalent binder and was not pursued further. Representative compounds from the CC*S and *CJS series were resynthesized off-DNA (Figures 4B-C), as both *cis* and *trans* isomers, for validation. None of these compounds produced appreciable CSPs against ^15^N-HuPrP90-231 using TROSY at 200 μM, suggesting that these hits were either false positives or have affinities too weak to be detected by NMR (Figures 4D-E).

**Figure 4.**
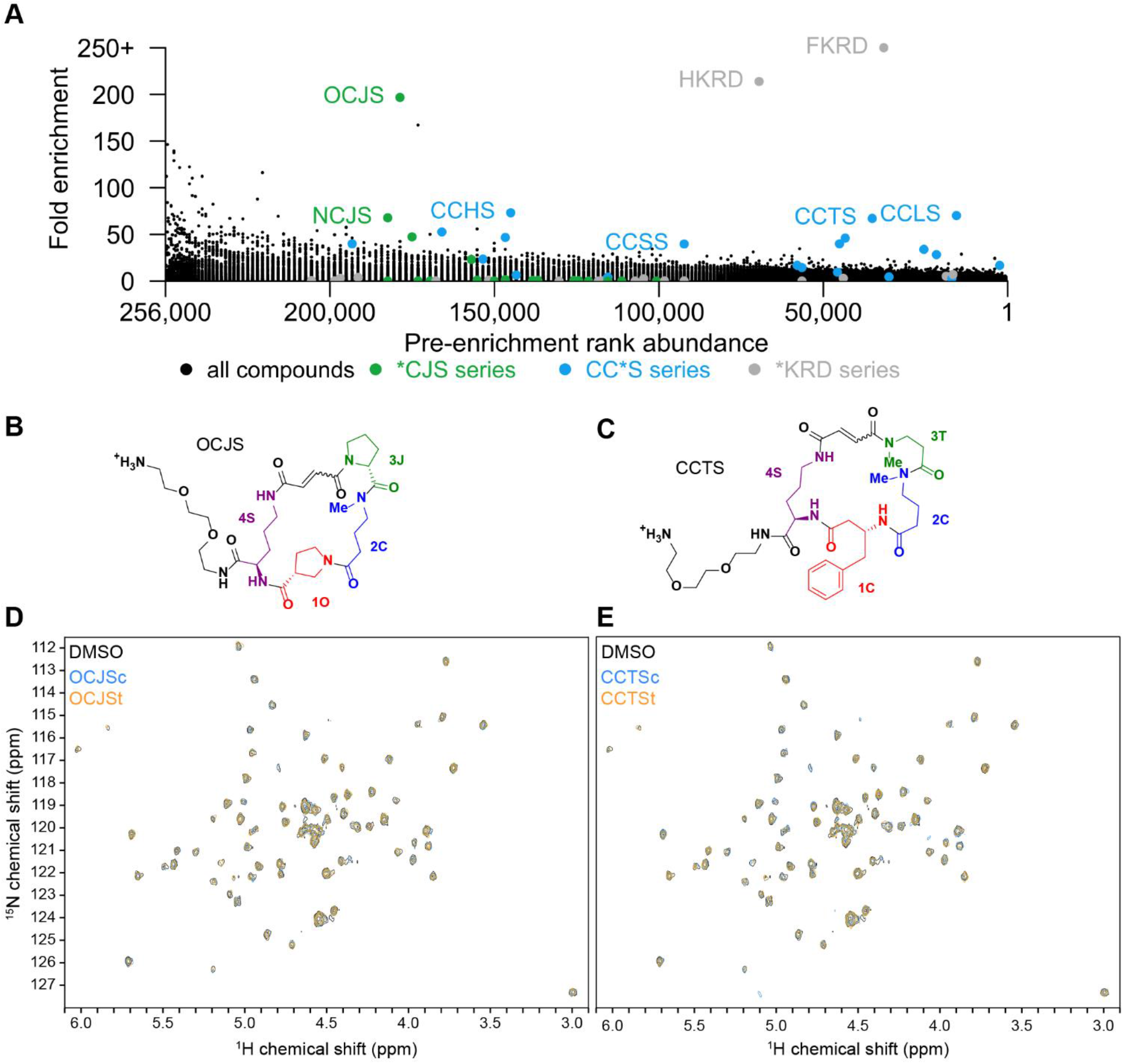
Selection of PrP binders from a DNA-encoded macrocycle library. A) Enrichment plot of macrocycle DEL compounds versus input rank. *KRD series compounds (grey) are frequent hitters. B and C) Structures of the DEL hits OCJS and CCTS synthesized off-DNA. DEL macrocycles are synthesized as stereoisomers on-DNA so each cis/trans isomer pair was synthesized off-DNA for testing (c, cis; t, trans). D and E) TROSY spectra of ^15^N-HuPrP90-231 with the cis and trans isomers (200 μM) of OJCS and CCTS.

### In silico screening

We used Atomwise’s AtomNet^®^ convolutional neural network method^58^ to search for compounds that bind PrP at a specific site. Since there are no reported structures of human PrP bound to a lead-like ligand, we instead used the reported structure of mouse PrP bound to promazine (PDB 4MA7)^65^ to create a homology model of human PrP also bound to promazine between helix 2 (α2) and the two beta strands (β1 and β2) (Figure 5A). The regions that were modeled share a high degree of sequence identity with only 12 amino acid differences over residues A117-R230 (mouse PrP numbering). The promazine binding site was screened against 6,922,894 molecules. After additional filtering, the top 81 compounds (Table S2) were selected as predicted binders and assayed for binding using both DSF and STD NMR. By DSF, none of the compounds (90 μM) increased the T_m_ of PrP more than three standard deviations (0.79 °C), and none of the compounds (100 μM) showed an appreciable STD signal in the presence of HuPrP90-231 (Figures 5B-C).

**Figure 5.**
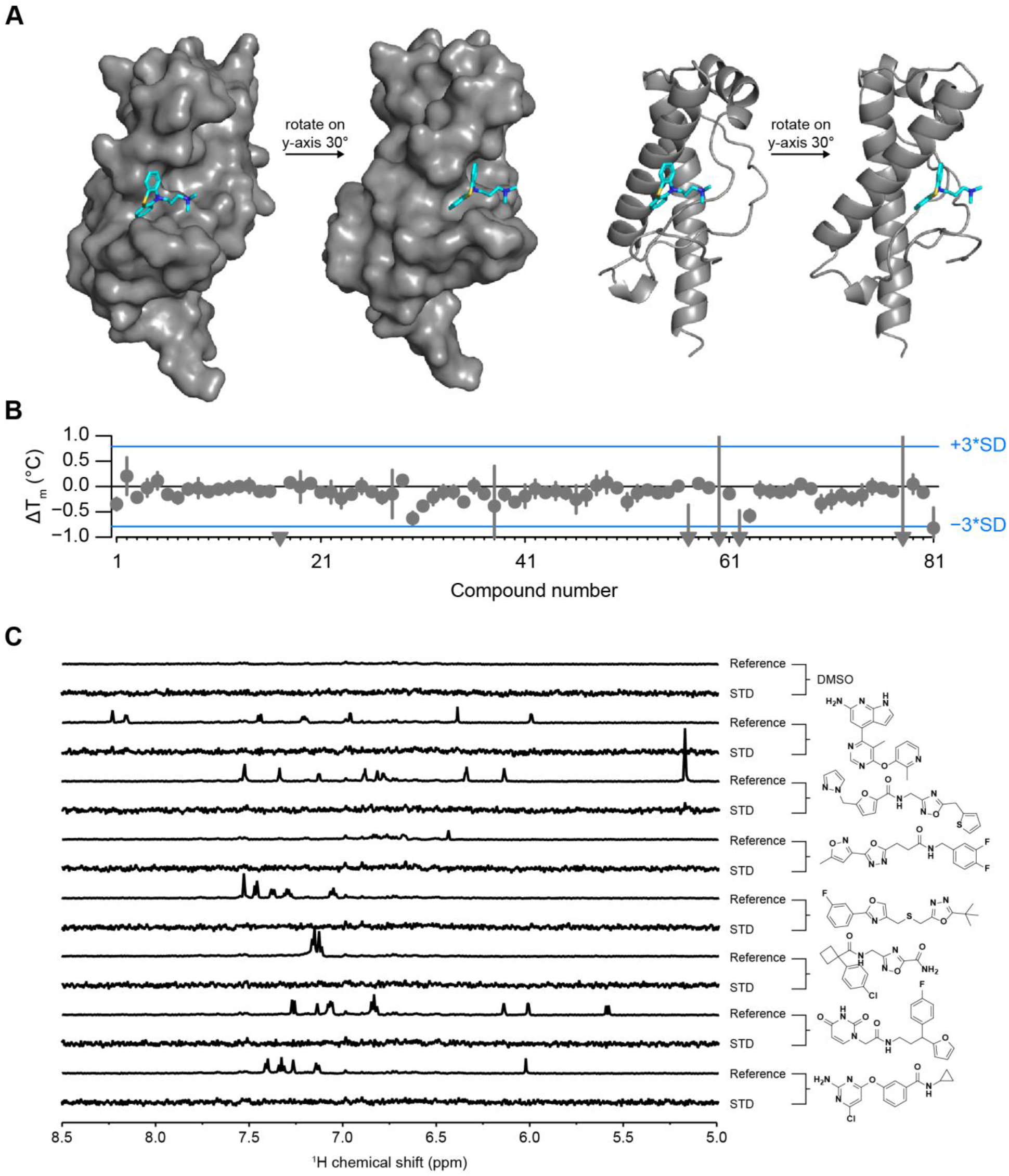
In silico screening. A) Homology model of HuPrP with ligand bound (promazine) that was used for neural network in silico screening. B) DSF ΔT_m_ values for all 81 Atomwise compounds with HuPrP90-231. Compounds that resulted in PrP shifts of less than one degree were plotted at −1. Error bars represent standard deviation of three measurements made on the same day using the same batch of protein-dye mix. SD of the DMSO conditions (n = 32 technical replicates per plate) was 0.26 °C. C) STD NMR of selected compounds that had a ΔT_m_ > 0.05 °C by DSF. STD spectra are scaled to 5X the reference spectrum intensity.

## DISCUSSION

We pursued four different screening modalities aimed at discovering binders of the human prion protein. Despite the large number of molecules tested and complementary approaches used, we were unable to identify any hits suitable for advancement into medicinal chemistry. Our fragment screening campaign identified compound **1** (5,6-dichloro-2-methyl-1*H*-benzimidazole) and several analogs that weakly bind PrP and were validated with orthogonal NMR assays. However, the poor affinity of these compounds (> 1 mM) coupled with the absence of improved binding of chemical analogs effectively precluded their validation through non-NMR methods, and none were pursued further. Meanwhile, our thermal shift, DNA-encoded library, and *in silico* screening approaches yielded no validated hits at all.

A variety of target-specific technical challenges may have contributed to our inability to identify binders by the approaches employed here. Some reports indicate that the transfer of NMR saturation is weak for smaller proteins, which may have produced false negatives in our STD NMR screens^66,67^. SYPRO Orange dye fluorescence in the presence of unfolded PrP was weak, which necessitated DSF screening at 30 μM protein concentration; compounds were accordingly screened at 100 μM, but solubility limitations may have prevented saturable binding with a maximum thermal shift. Our *in silico* screen utilized a homology model based on a crystal structure of promazine bound to mouse PrP, but promazine has not been shown to bind human PrP in solution, and promazine analogs that exert antiprion activity in cells appear to do so through an orthogonal mechanism^48^. In general, without a positive control available, it is difficult to guide the optimization of screening assays. Taken as a whole, our experimental screens cannot be considered definitive given their modest scale. But considering the diverse methods and compound sets employed, our results may hint toward relative rarity of PrP binders in chemical space.

Alternative screening approaches might also improve the probability of discovering a high affinity PrP binder. We used recombinantly expressed PrP from *E. coli* for our experiments, which lacks post translational modifications (PTMs) including two N-linked glycosylations and a GPI anchor. Purification of PrP from mammalian cells^68^ and insertion into nanodiscs^69^ or micelles might more faithfully recapitulate PrP’s PTMs and endogenous membrane environment, potentially yielding binding sites not present on recombinant PrP. Encouragingly, DEL screening has been used successfully with nanodisc immobilized proteins^70^. We could also extend our fragment-based drug design strategy by using chemoproteomics^71^ to directly assess PrP ligandability on the cell surface. Multiple approaches may be necessary, because targets with low NMR and thermal shift hit rates are reported to, on average, also have lower hit-to-lead development success rates^72^.

Overall, despite various technical limitations, our inability to identify even weak binders through multiple orthogonal screening modalities is striking. The absence of obvious binding pockets on PrP’s structure, together with the predominance of indirect mechanisms of action revealed in phenotypic screening campaigns, have led to the perception that PrP is a difficult target for small-molecule discovery^31^. Our data may provide some support for this conclusion. On balance, our results motivate an emphasis on non-small-molecule technologies, such as oligonucleotide therapeutics, as means for targeting PrP, but do not rule out the possibility that small-molecule binders could be discovered through an expanded screening effort.

## METHODS

### Log(P) and H-bond donor/acceptor calculations

SMILES strings were parsed to yield molecular weight, ALogP, and hydrogen bond donor and acceptor counts using RCDK^73^.

### Purification of HuPrP90-231 and HuPrP23-231

Recombinant PrP glycerol stocks were a generous gift from Byron Caughey and Andrew Hughson (NIAID Rocky Mountain Labs). The purification protocol was adapted from published procedures^74^. Two 4 mL cultures of *E. coli* were started from a glycerol stock in Terrific Broth (TB) with kanamycin (25 μg/mL) and chloramphenicol (25 μg/mL) and incubated (6 h, 37 °C, 220 rpm). Those cultures were used to inoculate a 1 L autoinduction media (AIM) (Millipore 71300) culture made with TB plus kanamycin (25 μg/mL) and chloramphenicol (25 μg/mL) in a 4 L baffled flask and incubated (22 h, 37 °C, 180 rpm). *E. coli* were harvested by centrifugation (4,300 *g*, 12 min, 4 °C) into four bottles (250 mL each) and the pellets were frozen at −80 °C. The following amounts of reagents used are based on a single pellet from 250 mL of media. The cell pellet was thawed at room temperature, resuspended in Lysis Buffer (14 mL) (Millipore 71456-4) by vortexing, and homogenized with a tissue homogenizer (60 s, 50% speed) using disposable tips. The homogenate was incubated (20 min, r.t., end-over-end agitation) and 20 μL of the whole-cell lysate was saved and diluted 1:20 (fraction “L”). The lysate was clarified by centrifugation (16,000 *g*, 20 min, 4 °C) and 20 μL of the supernatant was saved and diluted 1:20 (fraction “S”). Lysis Buffer (14 mL) was added to the pellet, which was homogenized with the tissue homogenizer (50% speed, 60 s) and incubated (20 min, r.t., end-over-end agitation). 0.1X Lysis Buffer (~20 mL) was added to a total volume of 34 mL, and the homogenate was centrifuged (16,000 *g*, 15 min, 4 °C). 30 μL of the supernatant was saved (fraction “W”). The pellet was resuspended in 30 mL 0.1X Lysis Buffer using a tissue homogenizer (50% speed, 1 min, r.t.) and centrifuged (16,000 *g*, 15 min, 4 °C). 30 μL of the supernatant (fraction “W2”) was saved. 10.5 mL of Unfolding Buffer (8 M guanidine HCl in 100 mM NaPO4 pH 8.0) was added to the inclusion body pellet and homogenized with the tissue homogenizer (50%-100% power, 1 min, r.t.). The homogenate was incubated (50 min, r.t., end-over-end agitation) and centrifuged (8,000 *g*, 5 min, 4 °C). 30 μL of the supernatant was saved (fraction “D”). All four supernatants were combined into a 50 mL conical tube and stored (7 days, 4 °C). 15 g of semi-dry Ni-NTA resin (Qiagen 30450) was weighed out into each of three conical tubes, denaturing buffer was added to 30 mL total volume, and the solution was incubated (10 min, end-over-end incubation, r.t.). Denatured PrP was evenly added to each of the three conical tubes and incubated (40 min, r.t.). PrP-bound resin was added to a column (Cytiva 28988948) and 30 μL of the unbound was saved (fraction “UB”). ÄKTA pure (Cytiva) lines were equilibrated with Denaturing Buffer (6 M Guanidinium HCl, 100 mM sodium phosphate pH 8.0) and Refolding Buffer (100 mM sodium phosphate, 10 mM Tris pH 8.0) and then 100% Denaturing Buffer. A gradient was run from 0-100% Refolding Buffer (2.25 mL/min, 240 min, 4 °C) and then 100% Refolding Buffer (2.25 mL/min, 30 min, 4 °C). The protein was eluted with a gradient from 0-100% Elution Buffer (500 mM imidazole, 100 mM sodium phosphate, pH 6.0) (6 mL/min, 45 min, 4 °C) and then 100% Elution Buffer (6 mL/min, 15 min, 4 °C). 30 μL of each fraction (fraction “#”) was saved. A 100 μL sample of the resin slurry was saved (fraction “B”). The fractions containing PrP were dialyzed (7 kDa MWCO, Thermo Fisher 68700) against 6 L of Dialysis Buffer (10 mM sodium phosphate pH 5.8) (overnight 4 °C) and 4 L of Dialysis Buffer (4-6 h, 4 °C). The dialyzed elution was centrifuged (4,300 *g*, 10 min, 4 °C) to pellet precipitated protein. A 30 μL sample of the final protein (fraction “F”) was saved. [HuPrP90-231] was measured by its absorbance at 280 nm (ε = 22,015 M^-1^ cm^-1^) (MW = 16.145 kDa). [HuPrP23-231] was measured by its absorbance at 280 nm (ε = 57,995 M^-1^ cm^-1^) (MW = 22.963 kDa). Protein was aliquotted, frozen in N2(l), and stored at −80 °C. For SDS-PAGE analysis, protein samples (30 μL) were mixed with 10 μL 4X Loading Buffer (4X LDS buffer [Thermo Fisher NP0007] with 10 mM TCEP [Thermo Fisher 77720]). For fraction “B” 25 μL of 4X Loading Buffer was added. Fractions D and UB were EtOH-precipitated before SDS-PAGE by adding 270 μL of EtOH, vortexing, and incubating on dry ice (5 min). D and UB were centrifuged (21,000 g, 5 min, 4 °C), and the supernatant was discarded. 300 μL 90% EtOH (−80 °C) was added, vortexed, and centrifuged (21,000 g, 5 min, 4 °C). The supernatant was discarded and the pellet was allowed to dry. 600 μL of 1X Loading Buffer was added to sample D and 40 μL 1X Loading Buffer was added to sample UB. All gel samples were incubated (90 °C, 5 min). Fractions were analyzed by SDS-PAGE with Coomassie staining (10 μL of each sample into a 15-well Bis-Tris NuPAGE gel (Thermo Fisher NP0321BOX), 180V, 40 min). The full amino acid sequences of HuPrP constructs are as follows. The N-terminal methionine of HuPrP90-231 was mostly removed by endogenous proteases as expected based on the second residue being glycine^75^; this was verified by intact protein LC-MS (Figure S5C). HuPrP23-231, however, retains the N-terminal methionine^76^, again as expected given the second residue is lysine^75^.

HuPrP90-231:

MGQGGGTHSQWNKPSKPKTNMKHMAGAAAAGAVVGGLGGYMLGSAMSRPIIHFGSDYEDRYYRENMHRYP NQVYYRPMDEYSNQNNFVHDCVNITIKQHTVTTTTKGENFTETDVKMMERVVEQMCITQYERESQAYYQRGSS

HuPrP23-231:

MKKRPKPGGWNTGGSRYPGQGSPGGNRYPPQGGGGWGQPHGGGWGQPHGGGWGQPHGGGWGQPHGGGWGQGGGTHSQWNKPSKPKTNMKHMAGAAAAGAVVGGLGGYMLGSAMSRPIIHFGSDYEDRYYRENMHRYPNQVYYRPMDEYSNQNNFVHDCVNITIKQHTVTTTTKGENFTETDVKMMERVVEQMCITQYERESQAYYQ RGSS

### Purification of ^15^N-HuPrP90-231 and ^15^N-HuPrP23-231

The same procedure was used to purify ^15^N-HuPrP with the following modifications. ^15^N AIM media was composed of 1X BioExpress ^15^N cell growth media (Cambridge isotope labs CGM-1000-N) in EMD Millipore Overnight Express induction system (Sigma-Aldrich 71300-M) with kanamycin (25 μg/mL) and chloramphenicol (25 μg/mL). Dialysis Buffer was 20 mM HEPES pH 7.4, 50 mM NaCl.

### NMR data acquisition and analysis

Spectra were acquired on a 600 MHz Bruker Avance III NMR Spectrometer equipped with a 5 mm QCI cryo-probe using 3 mm sample tubes (Bruker Z112272) and a SampleJet for sample handling. Spectra were analyzed in TopSpin version 4.0.2 and MestreNova version 10.0.1. Hit identification was performed by visual inspection of the data.

### ^19^F NMR screening

HuPrP23-231 (final concentration 9 μM) or matched dialysis buffer alone (20 mM HEPES pH 6.8, no-protein control condition) was combined with 10% D2O and 4.5% DMSO containing a pool of 10 ^19^F fragments per NMR tube (45 μM each) with a total volume of 200 μL for each sample. A ^19^F NMR spectrum was obtained for each sample using a standard ^1^H-decoupled one-pulse experiment with 64 scans and a spectral width of 237 ppm with the carrier frequency at −100 ppm. The sample temperature was 280 K. Fragment hits were identified by comparison of both the peak position and peak width between the control (no protein) sample and the protein-containing sample. For this screen minimal line-broadening was observed; fragment hits were identified by visual review of chemical shifts, with perturbation of ≥0.005 (3 Hz) as an approximate threshold. Hit peaks from pools were compared to reference spectra to identify the likely hit fragment. Resupplied fragments were tested by STD NMR and/or ^1^H-^15^N TROSY NMR as described below.

### STD NMR screening

HuPrP90-231 was buffer exchanged into 20 mM HEPES-d18 pH 7.4, 150 mM NaCl in ~99% D2O using a 5 kDa MWCO centrifugal concentrator (Millipore C7715). Pre-plated fragment pools were thawed in a dessicator at room temperature. For the Broad Institute 1^st^ and 2^nd^ generation STD libraries, 180 μL of HuPrP90-231 (11 μM) in deuterated buffer was added to each well, mixed with the fragments (1.6% DMSO-d6, v/v), and transferred immediately to a 3 mm NMR tube. The ChemBridge High Solubility Subset and Schreiber chiral fragment collection screens used HuPrP90-231 at 10 μM, 2 % DMSO-d6 (v/v), in 20 mM HEPES-d18 pH 6.8, 25 mM NaCl in ~99% D2O. Fragments were pooled with eight (Broad 1^st^ and 2^nd^ Gen STD and ChemBridge High Solubility Subset) or five (Schreiber chiral fragment collection) fragments per tube, always at a final concentration of 200 μM each. The Schreiber chiral fragment collection consisted of *n* = 381 compounds synthesized in-house, many of which have been described previously^77–79^. *In silico* screening hits (100 μM) were mixed with HuPrP90-231 (11 μM) in 20 mM HEPES-d18 pH 7.4, 150 mM NaCl in ~99% D_2_O with 1% DMSO-d6 (v/v). Ligand-observed screening was done using STD NMR. On-resonance irradiation of the protein was done at −0.25 ppm and off-resonance irradiation at 30 ppm. To saturate the protein, a 2 s train of 50 ms gaussian pulses separated by 1 ms delays was used. A 27 ms spin-lock pulse was used to suppress protein signals, and water suppression was accomplished using the excitation sculpting with gradients pulse scheme. The sample temperature was 280 K. Hit pools were identified by visual inspection and fragment hits were confirmed as singletons using the same experimental conditions as above. Compound **1** (5,6-dichloro-2-methyl-1*H*-benzimidazole) was purchased from two different vendors (Combi-Blocks HC-3145 and Key Organics PS-4319) and retested for an STD signal, which was reproducible across vendors. The compound from Key Organics was deemed more pure than other sources by NMR, thin-layer chromatography, and LC-MS and was used for the majority of experiments presented here.

### _1_H-^15^N TROSY NMR of ^15^N-HuPrP90-231 and ^15^N-HuPrP23-231

^15^N-HuPrP90-231 (50-60 μM) in 20 mM HEPES pH 7.4, 50 mM NaCl was combined with D2O (10%, v/v) and ligand (0-1 mM) or DMSO (2%, v/v), mixed and added to a 3 mm NMR tube. ^15^N-HuPrP23-231 in 20 mM HEPES pH 7.4, 150 mM NaCl was combined with D2O (10%, v/v) and compound **1** (0.3-1 mM) or DMSO (3%, v/v), mixed, and added to a 3 mm NMR tube. All concentrations are final. ^1^H-^15^N TROSY spectra were acquired at 298 K with 64 scans and 128 increments. Chemical shift perturbations (CSPs) were identified by visual inspection. Quantification of dose-response CSPs was performed with MestReNova version 10.0.1. Compound **1** was reproducible for CSPs across two different vendor sources used for retesting (described above).

### ^1^H-^15^N HSQC NMR of ^15^N-HuPrP90-231

^15^N-HuPrP90-231 (50 μM, 160 μL) in 20 mM HEPES pH 7.4, 50 mM NaCl was combined with 18 μL D2O (10%, v/v) and ligand (100 μM final concentration, 1.8 μL) or DMSO (1%, v/v), mixed and added to a 3 mm NMR tube. ^1^H-^15^N HSQC spectra were acquired using a 600 MHz Bruker Avance II spectrometer at 298 K. 2D data were processed and analyzed by using TopSpin version 4.0.2 software. CSPs were identified by visual inspection.

### AmpC aggregation counter-screen

Compounds were tested for colloidal aggregation using an AmpC β-lactamase counter-screen^62^. Recombinant *E. coli* AmpC was expressed in Rosetta cells and purified using a published protocol^80^. The enzymatic assay was performed in 50 mM sodium phosphate pH 7.0 ± 0.01% Triton X-100 (v/v) in clear UV-transparent 96-well half-area microplates (Corning 3679) in 150 μL final reaction volumes. Final concentration of DMSO was 1.0% (v/v). Compounds were incubated with 5 nM AmpC in 143.5 μL reaction solution for 5 min at r.t., followed by the addition of 1.5 μL of nitrocefin substrate (Cayman 15424) dissolved in DMSO (100 μM initial substrate concentration). Reaction solutions were gently mixed by multichannel pipette. Reaction progress was continuously monitored by absorbance at 482 nm for 5 min at r.t. on a SpectraMax M3 plate reader. Percent activity was calculated from reaction rates (slope) and normalized to DMSO-only controls after background subtraction with an enzyme-free reaction. Anacardic acid (AA), rottlerin, and 3’,3’’,5’,5’’-tetraiodophenolphthalein (TIPP) (Combi-Blocks QE-5474) were used as positive controls. The significance difference between detergent and non-detergent tests was defined as p < 0.01 (after correction for multiple comparisons). Four intra-run technical replicates were performed on same microplate.

### DSF screening

All concentrations are final assay concentrations unless otherwise indicated. All reagents were diluted in 20 mM HEPES pH 7.4, 150 mM NaCl, 1 mM EDTA. Protein was thawed and centrifuged (4500 *g*, 10 min, 4 °C) to pellet precipitate. HuPrP90-231 (30 μM) was mixed with SYPRO Orange dye (10X) (ThermoFisher S6651) and centrifuged (4500 *g*, 10 min, 4 °C) then decanted into a new bottle and covered in foil. Compound stocks dissolved in DMSO:H2O (90:10, v/v) were dispensed into 384-well barcoded plates (200 nL, 5 mM stock concentration, 100 μM final concentration) (4titude 4Ti-0381). 10 μL of protein-dye mix were added to each well with a Multidrop^™^ Combi Reagent Dispenser (ThermoFisher), shaken (2 min, r.t.), and centrifuged (1 min, r.t.). All Combi lines were covered to block ambient light. A Roche Lightcycler II was used for fluorescence measurements with filter sets 465/580. A temperature ramp from 30-90 °C, rate of 0.07 °C/s, and 6 acquisitions per second (6.5 min run) was used to collect the data. At the beginning of each day, 4 plates with no compounds were run to equilibrate the system. Melting temperatures (T_m_) were calculated by fitting the fluorescence data to a Boltzmann curve. ΔT_m_ values were calculated by taking the T_m_ with compound and subtracting it from the average of the DMSO control wells on each plate (*n* = 32 DMSO wells per plate). After removing error code flagged wells, positive ΔT_m_ hits were picked using the following criteria: ΔT_m_ > 3*MAD (0.17 °C), initial fluorescence intensity < 6. Negative ΔT_m_ hits were picked using the following criteria: −0.7 °C > ΔT_m_ > −9 °C, initial fluorescence intensity < 6. 1,129 hit compounds were tested in triplicate, all wells flagged with error codes 3 and 4 were removed, and hits were chosen based on the following criteria: ΔT_m_ sign must be the same as the primary screen, 19.93 > T_m_ > 0.3, initial fluorescence < 4, Δfluorescence of 2-17.1. Hits from the triplicate validation screen were filtered for frequent hitters and PAINS and 84 compounds were tested by HSQC NMR. DSF data were analyzed using Tibco Spotfire and RStudio.

### Differential scanning calorimetry

HuPrP90-231 (30 μM) was mixed with buffer or compound (100 μM) in 20 mM HEPES pH 7.4, 150 mM NaCl, 1 mM EDTA, 2% DMSO (v/v) (400 μL total volume per well). All concentrations are final assay concentrations. Data was acquired using MicroCal VP-Capillary DSC instrument from Malvern Panalytical and data was analyzed using Origin software provided by the vendor. A temperature ramp from 20-100 °C was conducted at a rate of 200 °C/h. Four buffer-only runs preceded the sample runs to equilibrate the system. *n* = 4 for DMSO controls and *n* = 1 for each compound. 3*SD was used as a hit cutoff.

### Macrocycle DEL screening

DNA-encoded library selection was performed as described using a 256,000-compound macrocycle library^55^. Briefly, 40 μg of HuPrP90-231, purified as described above, was loaded onto His Dynabeads (Invitrogen 10103D), relying on the intrinsic metal-binding properties of untagged PrP. Beads were washed, blocked, incubated with 50 μL of DNA-encoded library (60 min, 4 °C), washed three times, and protein was eluted with 300 mM imidazole. Barcodes were sequenced on an Illumina MiSeq and enrichment was calculated against a no protein condition run in parallel.

### In silico screening

The homology model of human PrP with bound ligand promazine was built on the template of the mouse PrP structure^65^ (PDB ID: 4MA7, chain A) using SWISS-MODEL^81^. The binding site is surrounded by residues V122, G124, L125, G126, Y128, Y162, I182, Q186, V189, T190 on the human PrP homology model.

The virtual screen was carried out using the AtomNet neural network, the first deep convolutional neural network for structure-based drug design^58,59^. A single global AtomNet model was employed to predict binding affinity of small molecules to a target protein. The model was trained with experimental *K*i, *K*d, and IC50 values of several million small molecules and protein structures spanning several thousand different proteins, curated from both public databases and proprietary sources. Because AtomNet is a global model, it can be applied to novel binding sites with no known ligands, a prerequisite to most target-specific machine-learning models. Another advantage of using a single global model in prospective predictions is that it helped prevent the so-called model overfitting. The following three-step procedure was applied to train the AtomNet model. The first step is to define the binding site on a given protein structure using a flooding algorithm^82^ based on an initial seed. The initial starting point of the flooding algorithm may be determined using either a bound ligand annotated in the PDB database or crucial residues as revealed by mutagenesis studies, or identification of catalytic motifs previously reported. The second step is to shift the coordinates of the protein-ligand co-complex to a three-dimensional Cartesian system with an origin at the center-of-mass of the binding site. Data augmentation was performed by randomly rotating and translating the protein structure around the center-of-mass of the binding site to prevent the neural network from memorizing a preferred orientation of the protein structure. The third step is to sample the conformations or poses of a small-molecule ligand within the binding site pocket. For a given ligand, an ensemble of poses was generated, and each of these poses represented a putative co-complex with the protein. Each generated co-complex was then rasterized into a fixed-size regular three-dimensional grid, where the values at each grid point represent the structural features that are present at each point. Similar to a photo pixel containing three separate channels representing the presence of red, green, and blue colors, our grid points represent the presence of different atom types. These grids serve as the input to a convolutional neural network and define the receptive field of the network. A network architecture of a 30×30×30 grid with 1Å spacing for the input layer, followed by five convolutional layers of 32×3^3^, 64×3^3^, 64×3^3^, 64×3^3^, 64×2^3^ (number of filters × filter-dimension), and a fully connected layer with 256 ReLU hidden units was used. The scores for each pose in the ensemble were combined through a weighted Boltzmann averaging to produce a final score. These scores were compared against the experimentally measured p*K*_i_ or pIC_50_ (converted from *K*_i_ or IC_50_) of the protein and ligand pair, and the weights of the neural network were adjusted to reduce the error between the predicted and experimentally measured affinity using a mean-square-error loss function. Training was done using the ADAM adaptive learning method^83^, the backpropagation algorithm, and mini-batches with 64 examples per gradient step.

The Mcule small-molecule library purchasable from the chemical vendor Mcule was used for the *in silico* screen. The original Mcule library version v20180817 containing approximately 10 million compounds in SMILES format was downloaded from Mcule’s website (https://mcule.com/). Every compound in the library was pushed through a standardization process including the removal of salts, isotopes and ions, and conversion to neutral form; conversion of functional groups and aromatic rings to consistent representations. Additional filters were applied on some molecular properties including molecular weight MW between 100 and 700 Daltons, total number of chiral centers in a molecule ≤ 6, total number of atoms in a molecule ≤ 60, total number of rotatable bonds ≤ 15, and only molecules containing C, N, S, H, O, P, B, halogens. Other filters such as toxicophores, Eli Lilly’s MedChem Rules^84^ and PAINS were also applied to remove compounds with undesirable substructures, resulting in the final library of 6,922,894 unique compounds.

For each small molecule, we generated a set of 64 poses within the binding site. Each of these poses was scored by the trained model, and the molecules were ranked by their scores. Due to a lack of a well-defined small molecule binding pocket on the human prion protein structure, there was low confidence in the predicted binders. Regardless, the top 50,000 ranking compounds were clustered based on chemical similarity and filtered for CNS drug-like properties using the Lipinski’s CNS rules^85^ with MW ≤ 400, clogP ≤ 5, number of hydrogen bond donors ≤ 3, and number of hydrogen bond acceptors ≤ 7. The final set of 81 compounds containing diverse chemical scaffolds were selected and sourced from Mcule.

### DSF of in silico hits and compound 1

Compounds were diluted to 1 mM in DMSO/HBS (20 mM HEPES pH 7.4, 150 mM NaCl) (20:80, v/v). 1 μL of compound was added to each well of a 384-well plate (90 μM final concentration). Mixtures of HuPrP90-231 (30 μM) were prepared with SYPRO Orange dye (10X) in 20 mM HEPES pH 7.4, 150 mM NaCl, 1 mM EDTA buffer then centrifuged (4000 *g*, 10 min, 4 °C) and decanted into a new tube. 10 μL were added to each compound and mixed (2% DMSO final concentration, v/v). A Roche Lightcycler II was used for fluorescence measurements with filter sets 465/580. A temperature ramp from 30-90 °C, rate of 0.07 °C/s, and 8 acquisitions per second (~18 min per plate) was used to collect the data. Three independent plates (using the same protein-dye mix) were measured with *n* = 32 for DMSO controls per plate and *n* = 1 for each compound per plate. ΔT_m_ values were calculated by subtracting the average apo T_m_ from the T_m_ with compound. Three ΔT_m_ values per compound were averaged and plotted. Error bars represent standard deviation (SD = 0.26 °C). For DSF of compound **1** with HuPrP90-231, the same experimental parameters were used as above except compound **1** was used at 500 μM concentration and *n* = 8 intra-run technical replicates performed on the same assay plate.

### Intact protein LC-MS

Purified HuPrP90-231 was diluted to 2 μM in 20 mM HEPES pH 7.4, 50 mM NaCl. 1 μL of diluted protein was injected onto a Waters BioAccord LC-ToF (composed of an ACQUITY I-Class UPLC and RDa detector with ESI source). Mobile phase A consisted of 0.1% formic acid (Millipore LiChroPur) in LC-MS grade water (JTBaker) and mobile phase B consisted of 0.1% formic acid in LC-MS grade acetonitrile (JTBaker). Protein was trapped on a C4 column (ACQUITY UPLC Protein BEH, 300Å, 1.7 μm, 2.1 × 50 mm) held at 80 °C for the entire analysis. The protein was desalted for one minute before elution with a gradient of 5% to 85% mobile phase B in 2.5 min. Ionization was performed with 55 V cone voltage and 550 °C ionization temperature. The instrument scan rate was 0.2 scans/s over 50 to 2000 m/z. PrP eluted at an observed retention time of 2 min. The PrP charge envelope was deconvoluted into the intact mass using the MaxEnt1 function using UNIFI software (Waters).

## Code and data availability

Raw data and source code will be made available in a public GitHub repository: http://github.com/ericminikel/binder_screening

## AUTHOR CONTRIBUTIONS

AGR, MFM, SMV, AJL, KTN, DSA, SLS, EVM, conceived and designed the study. AGR, MFM, DC, SMV, JLD, AJL, AIC, DLU, JBY, RNS, OKS, JRS, VLR, JAM, MS, MCN, KTN, and EVM performed the experiments and interpreted data. MFM, SMV, AJL, CTL, AJC, MCN, KTN, SKW, DRL, FFW, VKK, DSA, SLS, and EVM supervised the research. AGR and EVM wrote the paper. All authors reviewed and edited the paper.

## ACKNOWLEDGMENTS

This work was supported by the National Institutes of Health (F31 AI122592 to EVM, R35 GM118062 to DRL supporting AIC and DLU, and T32 HL007627 to JLD), Prion Alliance, BroadIgnite, and an anonymous organization. Atomwise provided effort and reagents in kind as part of an Artificial Intelligence Molecular Screen (AIMS) Award. Novartis provided reagents, expertise in screening sciences and efforts to perform the work as part of a research collaboration agreement. We thank Dr. Kirk Clark (Novartis) for guidance on the DSF screen and related NMR validation experiments.

## SUPPLEMENT

### Supplemental Tables (see Excel files)

**Supplemental Table 1.** All compound **1** analogs tested for HuPrP90-231 binding. +, positive STD/TROSY signal; -, no STD/TROSY signal; ±, borderline positive STD/TROSY signal. Blank indicates not tested. All spectra were assessed by visual inspection. For precipitation notes, “+” indicates that precipitate was observed during the experiment, and “±” indicates mild or questionable precipitation. R_1-4_ and X groups correspond to the molecule in Table 2.

**Supplemental Table 2.** *In silico* screening hits tested by DSF and STD NMR.

### Supplemental Figures

**Supplemental Figure 1.**
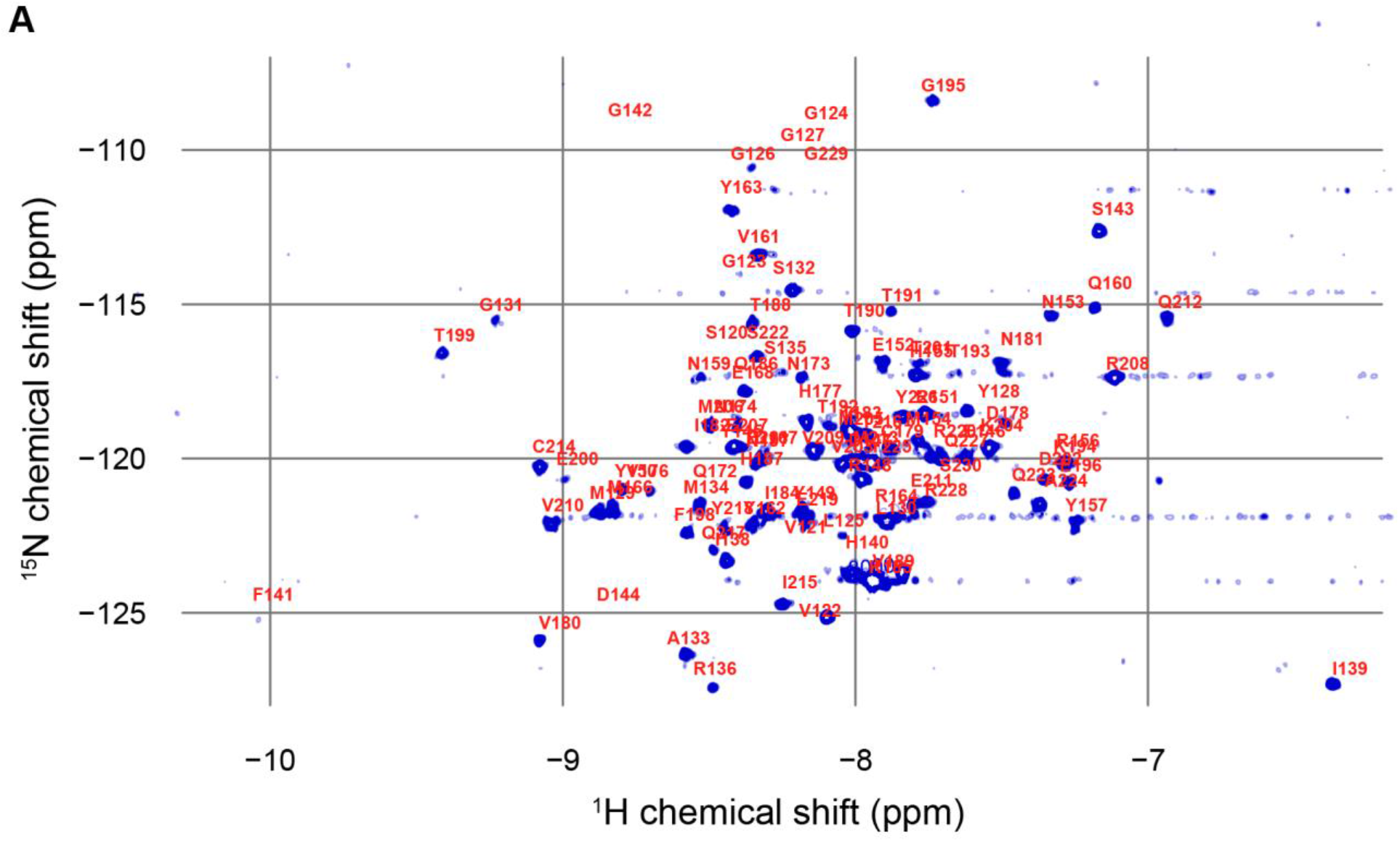
A) TROSY spectrum of ^15^N-HuPrP90-231 overlaid with residue annotations from BMRB #5713^61^.

**Supplemental Figure 2.**
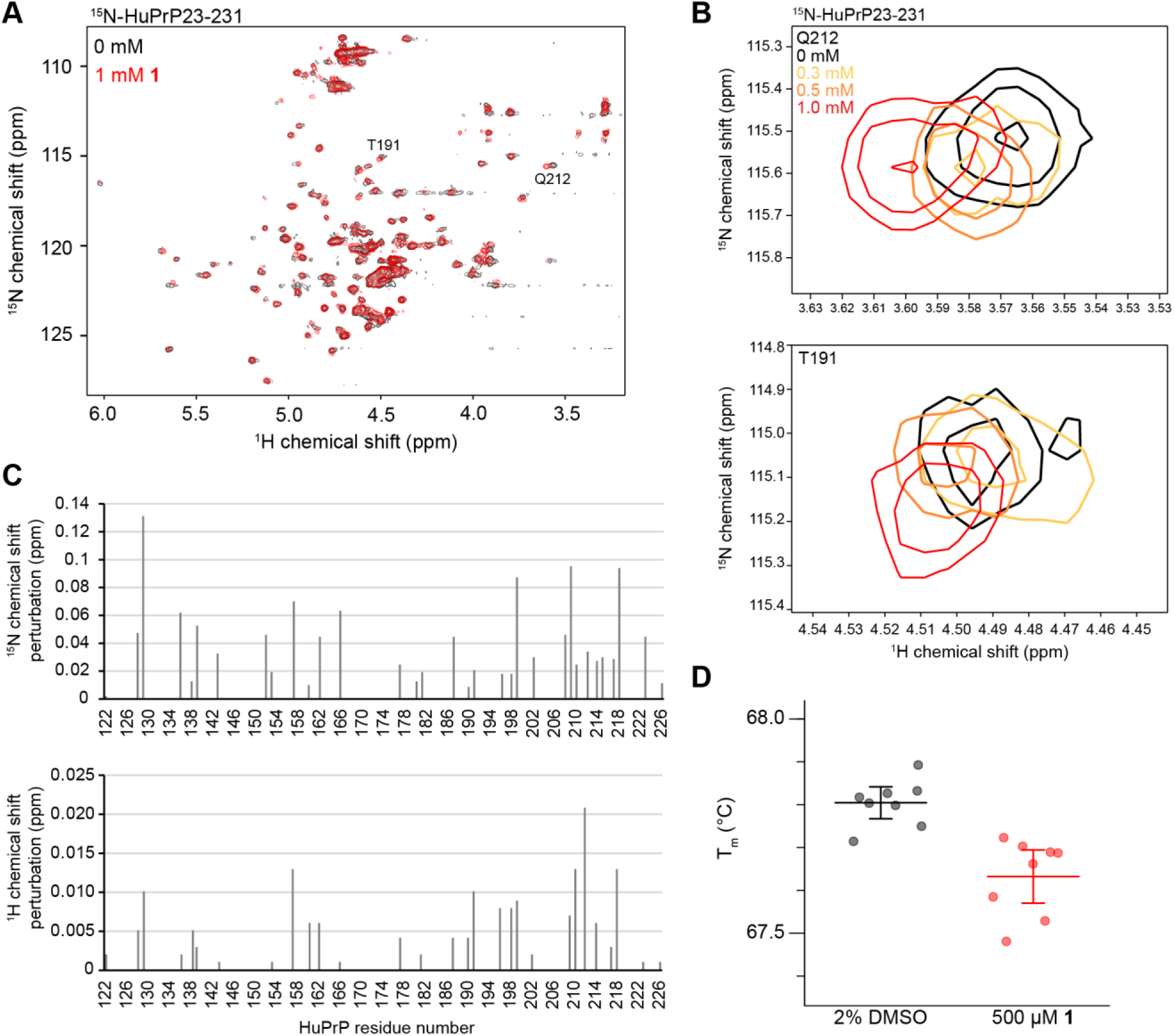
A) TROSY NMR of ^15^N-HuPrP23-231 with DMSO (black) or 1 mM **1** (red). B) Concentration dependent CSPs of residues Q212 and T191 upon addition of **1**. C) CSPs for ^15^N and ^1^H from ^15^N-HuPrP90-231 with 0.75 mM **1** before normalization as in Figure 2C. D) DSF T_m_ values for HuPrP90-231 with and without 500 μM **1**. Error bars represent 95% CI of *n* = 8 intra-run technical replicates performed on the same assay plate.

**Supplemental Figure 3.**
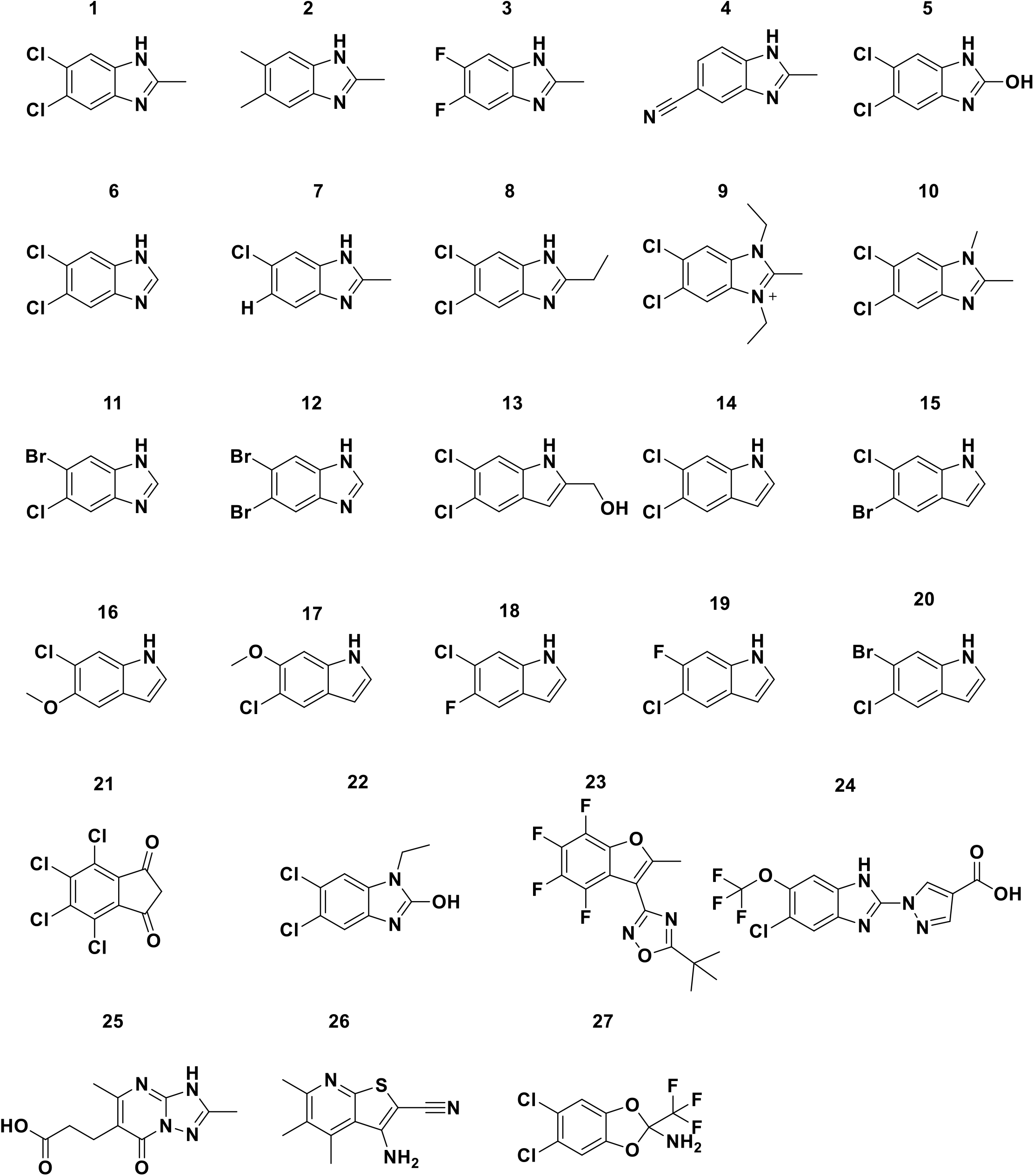
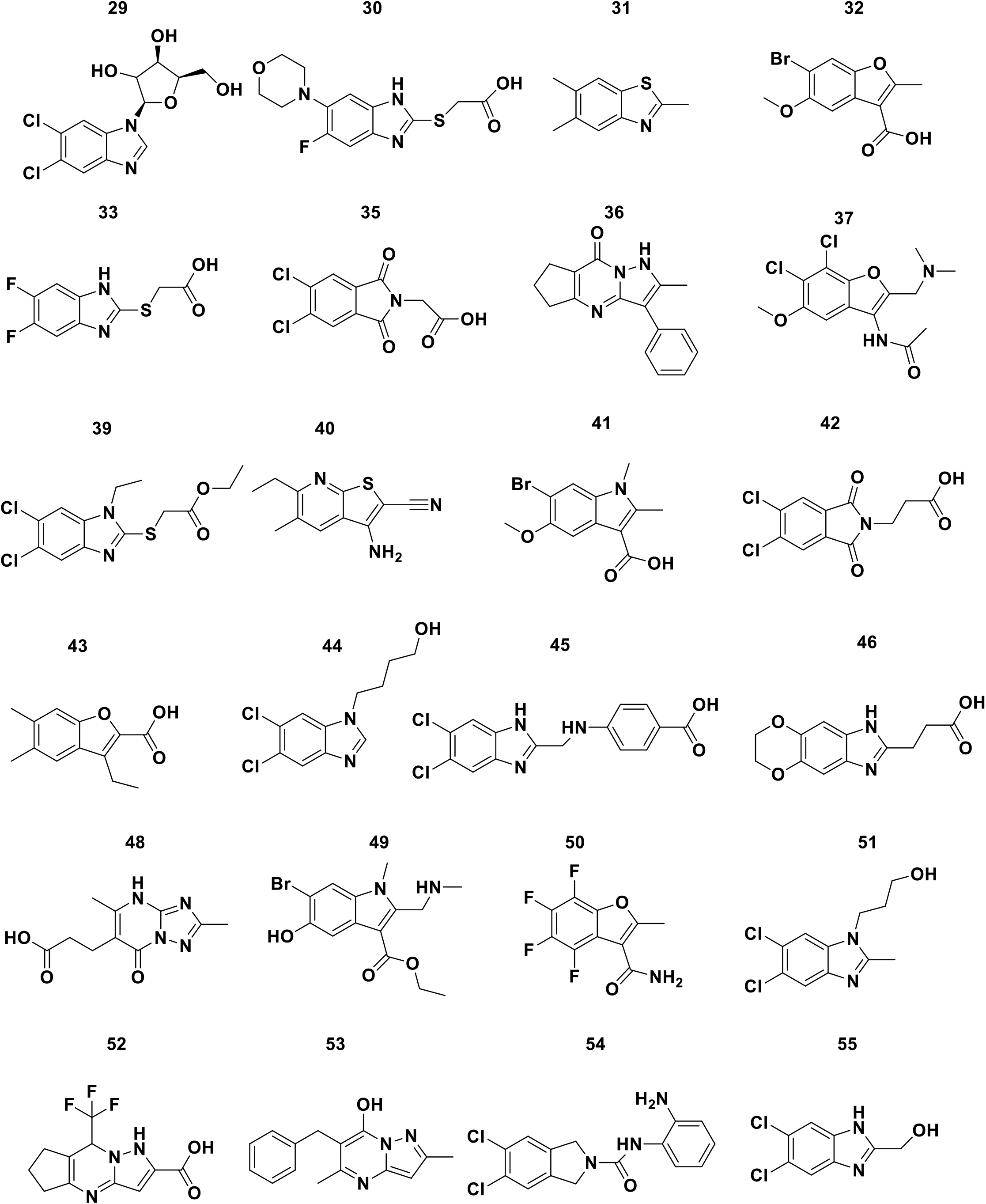
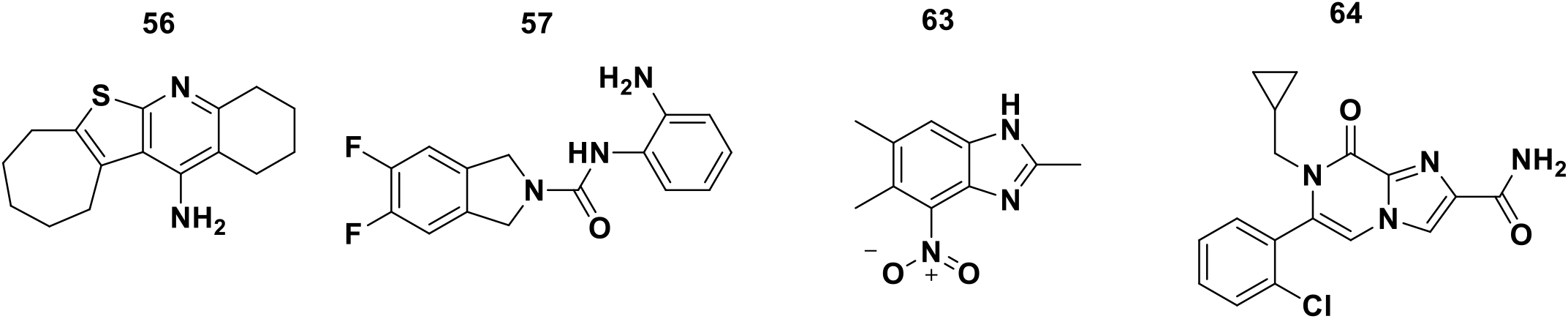
Analogs of **1** tested by STD NMR and TROSY NMR. Numbers correspond to compound number in Table 2 and Table S1.

**Supplemental Figure 4.**
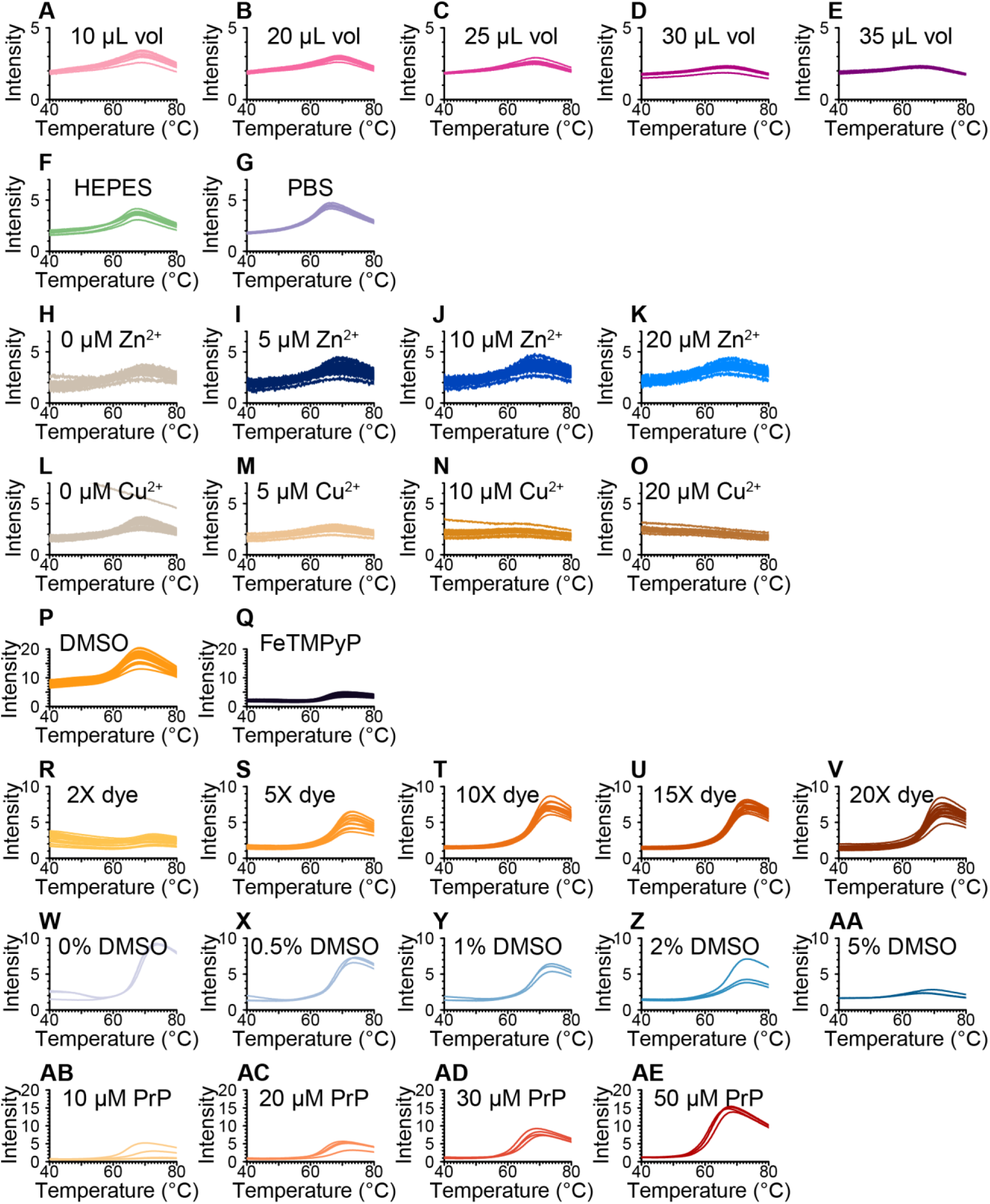
Selected conditions explored during the DSF assay development process. Assay development was initially undertaken at the Broad Institute **(A-Q)** before the assay was transferred to Novartis **(R-AE)**. **A-E)** Assay volume. Because a prior report^31^ used 150 μL assay volume in a 96-well format, we asked whether signal would be improved by increasing assay volume within the limit allowed by our 40 μL 384-well plates. HuPrP23-231 (5.7 μM) with SYPRO Orange (8.75X) and 1% DMSO (v/v) in 20 mM HEPES pH 6.8, 25 mM NaCl, 1 mM EDTA (hereafter HEPES Buffer). Because no improvement in signal was observed, 10 μL volume is used in subsequent experiments. **F-G)** Buffer. HuPrP23-231 (10 μM) with SYPRO Orange (15X) and 1% DMSO in either HEPES Buffer or 137 mM NaCl, 2.7 mM KCl, 10 mM Na2PO4, 1.8 mM KH2PO4 pH 7.4 (CSH PBS; doi:10.1101/pdb.rec8247). **H-K)** Zinc titration. HuPrP23-231 (5 μM) with SYPRO Orange (15X) and 1% DMSO (v/v) in CSH PBS with 0, 1, 2, or 4 molar equivalents of ZnSO_4_. **L-O)** Copper titration. HuPrP23-231 (5 μM) with SYPRO Orange 15X and 1% DMSO in CSH PBS with 0, 1, 2, or 4 equivalents of CuSO_4_. **P-Q)** Candidate positive control. HuPrP23-231 (50 μM) with SYPRO Orange (10X) and 2% DMSO (v/v) in HEPES Buffer, with or without 20 μM of the iron porphyrin FeTMPyP (Fe(III)tetrakis (1-methyl-4-pyridyl) porphyrin pentachloride; CAS #133314-07-5; Cayman Chemical #75854). Note that EDTA may chelate the iron from TMPyP, and unmetallated porphyrins are reported to be less active than metallated ones^37^; moreover, because FeTMPyP is substoichiometric to PrP, even if it binds with a *K*_d_ of ~1 μM as reported^46^, only a minority of PrP will be bound. For both of these reasons, the dramatic change in melting curve observed in this experiment is likely an artifact of FeTMPyP fluorescence. **R-V)** Dye titration. 30 μM HuPrP90-231 with varying SYPRO Orange concentrations. **W-AA)** DMSO titration. HuPrP90-231 (30 μM) with SYPRO Orange (10X) and varying DMSO concentrations (v/v) in 20 mM HEPES pH 7.5, 150 mM NaCl. **AB-AE)** Protein titration. HuPrP90-231 at varying concentrations with SYPRO Orange (10X) in 20 mM HEPES pH 7.0, 150 mM NaCl.

**Supplemental Figure 5.**
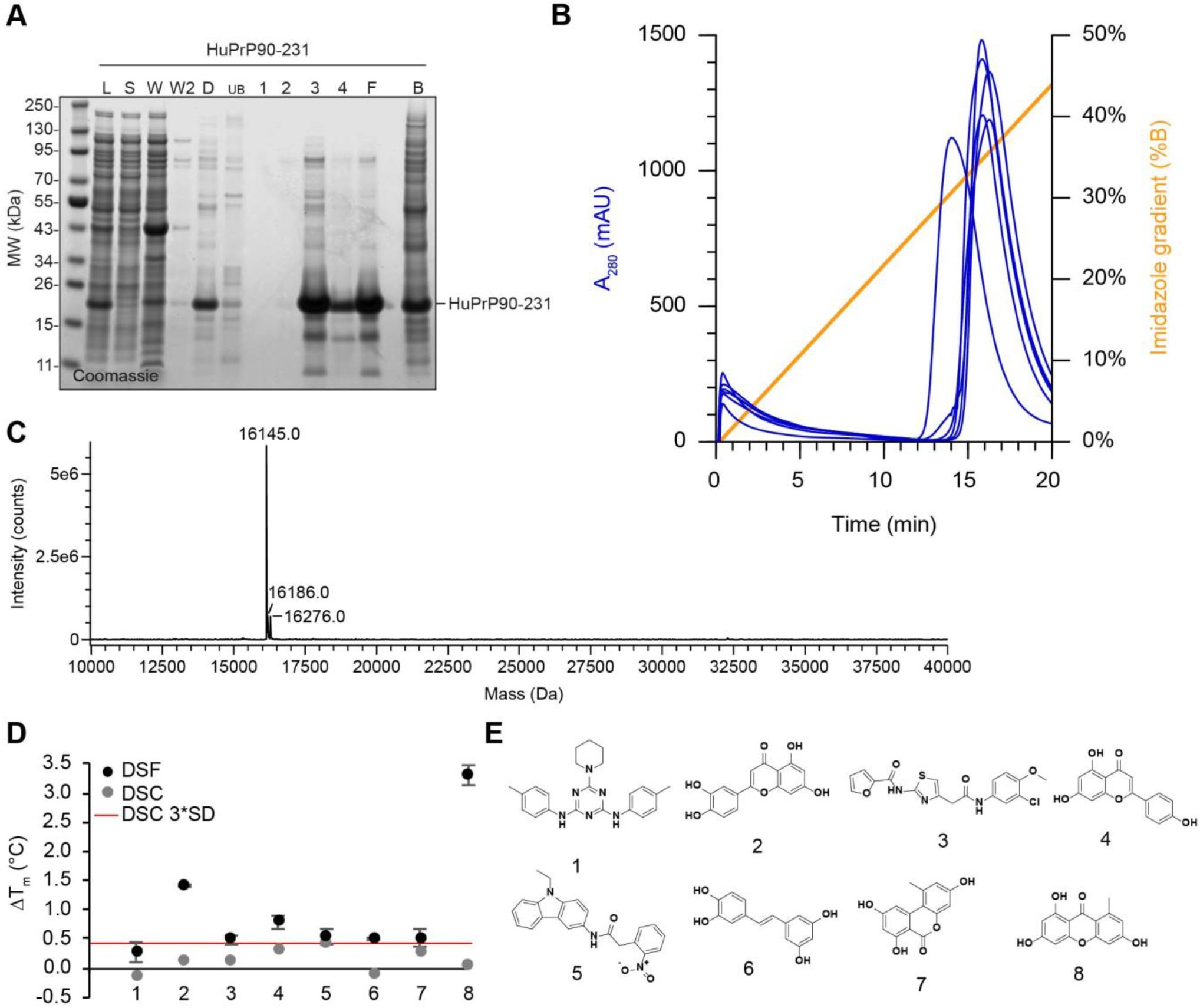
A) Example SDS-PAGE of HuPrP90-231 protein purification used for the DSF screen. L, whole-cell lysate (diluted 1:20); S, soluble fraction (diluted 1:20); W, first inclusion body wash; W2, second inclusion body wash; D, denatured protein (diluted 1:20) post centrifugation; UB, unbound; 1-4, AKTA elution fractions; B, Ni-NTA resin after elution; F, final PrP sample stored at −80 °C. B) AKTA elution traces for six separate purifications of HuPrP90-231, which elutes between 12-20 minutes. C) Deconvoluted charge envelope of HuPrP90-231 from intact protein LC-MS. D) Differential scanning calorimetry of eight (*n* = 1) selected positive shifters present in the Broad Institute’s compound collection along with corresponding DSF data from the validation screen. 3*SD = 0.41 °C for *n* = 4 apo (DMSO) replicates. E) Compound structures corresponding to panel D.

